# Flower size evolution in the Southwest Pacific

**DOI:** 10.1101/2023.12.12.571326

**Authors:** Riccardo Ciarle, Kevin C. Burns, Fabio Mologni

**Affiliations:** Te Kura Mātauranga Koiora | School of Biological Sciences, Te Herenga Waka | Victoria University of Wellington, PO Box 600, Wellington, Aotearoa New Zealand; Division of BioInvasions, Global Change & Macroecology, University of Vienna, Austria

**Keywords:** Island rule, flower size, island biogeography, plant evolution, Southwest Pacific, Island syndrome

## Abstract

**Background and Aims:** Despite accelerating interest in island evolution, the general evolutionary trajectories of island flowers remain poorly understood. In particular the island rule, which posits that small organisms become larger and large organisms to become smaller after island colonization, while tested in various plant traits, has never been tested in flower size. Here, we provide the first test for the island rule in flower size for animal- and wind-pollinated flowers, and the first evidence for generalized *in-situ* evolution of flower size on islands.

**Methods:** Focusing on 10 archipelagos in the Southwest Pacific, we amassed a dataset comprising 129 independent colonization events, by pairing each island endemic to its closest mainland relative. We then tested for the island rule in flower size and for gigantism/dwarfism in floral display for animal- and wind-pollinated flowers.

**Key results:** Animal-pollinated flowers followed the island rule, while wind-pollinated flowers did not, instead showing evidence of gigantism. Results remained consistent after controlling for breeding system, mainland source pool, degree of taxonomic differentiation, taxonomic family, and island type.

**Conclusions:** While *in situ* evolution of flower size is widespread on islands in the Southwest Pacific, animal- and wind-pollinated flowers exhibited unexpected and markedly different evolutionary trajectories. Further studies are needed to understand the mechanisms behind these patterns.

## Introduction

Island flowers often differ from their mainland relatives, being typically dull-coloured and inconspicuous (Whittaker et al., 2023). They are often self-compatible and possess generalized morphologies that allow pollination by a wide array of pollinators (Baker, 1967; Carlquist, 1974; Barrett, 1996; Crawford et al., 2011; Burns, 2019). These differences suggest the existence of an island floral syndrome (i.e. a set of predictable differences between island and mainland flowers) which is still largely unexplored (Whittaker et al., 2023).

The island floral syndrome can arise from evolutionary processes and/or from insular assembly rules (i.e. patterns arising from biased colonization/establishment and not in-situ evolution, *sensu* Whittaker et al., 2023). At the moment, most components of the island floral syndrome are regarded as insular assembly rules (but see Hetherington-Rauth and Johnson, 2020). For example, plants capable of self-fertilization are overrepresented on islands as they can establish new populations more readily (Baker, 1967; Pannel et al., 2015; Grossenbacher et al., 2017; Burns, 2019). In turn, self-compatible plants typically have smaller flowers (Goodwillie et al., 2010). Whether components of the island floral syndrome can arise from in-situ evolutionary processes is still unclear (Bernardello et al., 2001; Newstrom and Robertson, 2005; Burns, 2019).

Directional evolutionary processes in island species traits can result either in overall trends toward gigantism or dwarfism (Burns et al., 2012; Kavanagh and Burns, 2014), or in the ‘island rule’, a pattern in which, after island colonization, small species become larger and large species to become smaller. While it remains one of the most controversial topics in biogeography (Van Valen, 1973; Lomolino, 2005; Meiri et al., 2008; Lomolino et al., 2012; Faurby and Svenning, 2016; Itescu et al., 2018; Rebouças et al., 2018; Lokatis and Jeschke, 2018; Benítez-López et al., 2021), evidence for the island rule has recently been found for plant stature and leaf area on islands in the Southwest Pacific (Biddick et al., 2019). Yet, whether other plant traits conform to an island-rule-like pattern remains unclear.

Islands are known for their paucity of pollinators (Burns, 2019; Whittaker et al., 2023). Changes in pollinator abundance between islands and the mainland could generate directional selective pressures in floral display (i.e. flower size ⋅ number of flowers per individual), causing animal-pollinated floral displays to increase/decrease and wind-pollinated floral displays to remain unchanged (Parachnowitsch and Kessler, 2010; Caruso et al., 2019; Brunet et al., 2021). Pollen limitation could intensify selection for traits promoting outcrossing through increased floral display, potentially leading to larger flowers (Harder et al., 2010; Thomann et al., 2013; Panique and Caruso, 2020). Alternatively, pollinator decline and mate limitation could select for selfing mechanisms, leading to smaller floral displays and smaller flowers in animal-pollinated species (Elle and Carney, 2003; Thomann et al., 2013).

Flower size of both animal- and wind-pollinated species could alternatively follow the island rule as a by-product of selection acting on flower-correlated traits. In particular, leaf area and seed size are known to covary with sepal and corolla size and were shown to evolve predictably on islands (Niklas, 1994; Burns et al., 2012; Biddick et al., 2019; E-Vojtkó et al., 2022; Whittaker et al., 2023). As such, flower size might not be under direct selection, but follow the island rule because of its allometric relationships with leaf area and seed size. Under this hypothesis, both animal- and wind-pollinated island flowers should differ from their mainland relatives, and these patterns should mirror those of leaf area and seed size.

Different plant lineages can respond independently to insular selective pressures. For example, the genetic, morphological and evolutionary history of certain taxonomic families make them more likely to evolve generalized floral structures or to break down mechanisms of self-incompatibility (Goldberg et al., 2010; Minelli, 2018; Cossard et al., 2021). Similarly, plants with different breeding systems are typically characterized by flowers of different size and colour (Krizek and Anderson, 2013; Salisbury et al., 2017), rely on different pollination strategies and typically respond to changes in pollinator pressures in different ways (Barrett, 1998). Additionally, evolutionary pressures will vary depending on the abiotic and biotic characteristics of the islands themselves (MacArthur and Wilson, 1967; Whittaker et al., 2023). Consequently, to test for general patterns of flower size evolution on islands, it is vital to include multiple plant lineages with different pollination modes, evolutionary histories and breeding systems that occur across multiple archipelagos.

In this study, we used a phylogenetic comparative approach to test whether island animal-pollinated and wind-pollinated flowers evolve toward gigantism/dwarfism or follow the island rule. Specifically, using a newly derived dataset representing 129 independent colonization events on 10 archipelagos in the Southwest Pacific, we asked two questions: I) Does floral display of animal-pollinated species increase/decrease, while floral display of wind-pollinated species remain unchanged? II) Do both animal-pollinated and wind-pollinated flowers evolve to follow the island rule, and does this pattern mirror those of leaf area and seed size?

## Methods

### Study area

We focused on the endemic flora of 10 archipelagos distributed across 25° of latitude (29°04’ S -54°37’ S) around New Zealand (Fig 1). The island groups can roughly be divided into the sub-tropical archipelagos of Lord Howe, Norfolk, Kermadec, and Three Kings, and the cool-temperate/subantarctic archipelagos of Chatham, Snares, Antipodes, Auckland, Campbell and Macquarie. All island systems are oceanic in origin, with faunas and floras derived by overwater dispersal from New Zealand and Australia or vicariance (Chown et al., 1998; Rutledge et al., 2003; Pendlebury and Barnes-Keoghan, 2007; Lord, 2015; Fitzgerald, 2020). We restricted analyses to these island systems so that our set of island taxa would be derived from a few well-known mainland floras, sharing similar ecological, geological and evolutionary histories. Chatham, Snares, Antipodes, Auckland, Campbell, Three Kings, Kermadec and Macquarie were included as most of their endemic species originated from the New Zealand mainland. Lord Howe and Norfolk islands were included as most of their endemic species originated from Australia. We included endemics from 2 mainland sources to increase the number of taxonomic families present and to account for the possibility that endemics originating from very different mainland source pools (i.e. New Zealand and Australia) would respond differently to insular environments.

**Figure 1.**
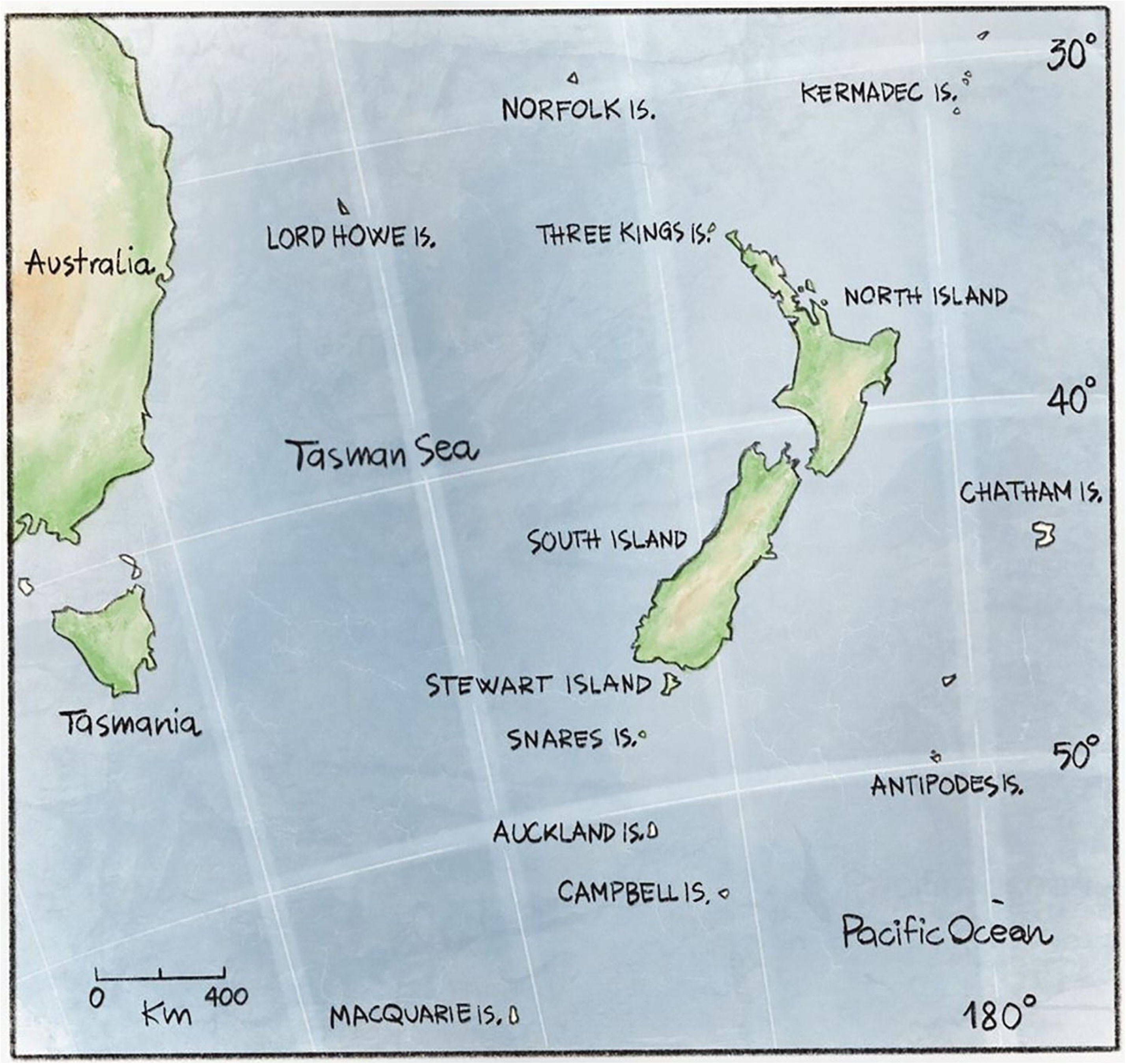
Map of the studied archipelagos.

### Species selection

To test flower size evolution in animal- and wind-pollinated species, we compiled a list of island/mainland pairs of taxa. Each mainland taxon is the sister group of the island taxon so that each pair represents a phylogenetically independent comparison.

We first compiled a list of endemic species and subspecies on each archipelago using the most recent data available (Oliver, 1948; Godley, 1969; Taylor, 1971; Johnson and Campbell, 1975; Meurk, 1977; Sykes, 1977; Godley, 1989; Wright, 1983; Green, 1994; Meurk et al., 1994; Sykes and West, 1996; Amey et al., 2007; De Lange et al., 2011; Lord et al., 2013; De Lange, 2014; De Salas and Baker, 2015). Because botanical sources varied widely in age, all species names were cross-checked for synonymy, keeping the New Zealand Plant Conservation Network (2023) and the Atlas of Living Australia (2023) as taxonomic references.

We then identified the mainland sister taxon of each island taxon using published molecular phylogenies. We searched Google Scholar for phylogenies using the species or genus name and the Boolean operator “AND” along the term “phylogen*” (e.g. *Coprosma* AND phylogen*). When multiple phylogenies were available, we selected the phylogeny that used the most comprehensive genetic dataset, used either Bayesian inference or Maximum likelihood for phylogenetic reconstruction, and showed the strongest support at each node (Supplementary 1). For each phylogeny, we included all representatives of a given genus on the mainland to ensure that any possible sister taxon of the island endemic was considered. Species names were standardized following the New Zealand Plant Conservation Network (2023) and Atlas of Living Australia (2023). When molecular phylogenies were unavailable or did not include all mainland species within a genus, sister taxa were identified based on morphological evidence. To do this, we focused on vegetative traits such as leaf and stem morphology (e.g. leaf lamina size and shape, venation structure, petiole length). As reproductive traits represent the dependent variable of our analyses, any reproductive trait was excluded for identification. As without phylogenies we could not identify the closest relative with certainty, multiple mainland species within the genus were selected and their floral traits averaged. If the genus comprised less than five species on the mainland (e.g. genus *Streblus*), all species were included as mainland relatives. If the genus comprised more than five species on the mainland (e.g. genus *Exocarpos*), we included as mainland relatives up to five species based on morphological affinities. (Supplementary 2). We excluded island taxa if mainland taxa could not be identified using either phylogeny or taxonomy.

### Trait measurements

Each island/mainland pair was categorized as animal-pollinated or wind-pollinated, and as monomorphic (including hermaphroditic, monoecious, andromonoecious, and gynomonoecious species) or dimorphic (including dioecious and gynodioecious species) using McGlone and Richardson (2023), Atlas of Living Australia (2023), Flora of New South Wales Online (2023) and the New Zealand Plant Conservation Network (2023).

To test our first hypothesis, floral display was defined as flower number ⋅ flower size. For flower number, we counted the number of flowers per individual. When unavailable, we used the number of flowers per inflorescence as an estimate of floral display. This can be a reliable estimate as inflorescences typically behave as autonomous modules among which resources cannot be reallocated (Herrera, 1991; Torices and Méndez, 2014; Dai et al., 2018).

To test our first and second hypotheses, flower size of animal-pollinated flowers was measured using the unit of pollinator attraction (Faegri and Van der Pijl, 1979), i.e. the floral trait bearing the primary attractiveness function (e.g. petals), so that each trait had a comparable biological function across families. For radially symmetric flowers with fused petals and petal lobes (e.g. Plantaginaceae, genus *Veronica*), we characterized lobe length, lobe width, or corolla diameter. Tube length was collected but not included in the analysis as it is not a measure of advertisement but of accessibility (Dafni, 1992). For radially symmetric flowers with free petals, we characterized petal length, petal width or corolla diameter. For zygomorphic flowers, we used the length and width of the organs most involved in advertisement (e.g. banner and wing of Fabaceae flowers). For the Myrtaceae family, we characterized both stamen length and petal length, as in this family the stamens are often involved in pollinator attraction (Vasconcelos et al., 2019). For Asteraceae, we collected ray floret length, disk diameter or capitulum diameter. For dimorphic taxa, we always used measurements from male flowers, as we expect the male perianth to be the one under stronger selective pressures (Barbot et al., 2023).

Flower size of wind-pollinated species was measured the sterile parts of flowers, as these traits are known to correlate with leaf area and seed size (Burns et al., 2012; Biddick et al., 2019; E-Vojtkó et al., 2022). For radially symmetric flowers with fused petals (e.g. Plantaginaceae, genus *Plantago*), we characterized lobe length, lobe width, or corolla diameter. For radially symmetric flowers with free petals (e.g. Arecaceae) we characterized petal length or corolla diameter. For flowers with undifferentiated sepals and petals (e.g. Juncaceae), we used tepal length. For Poaceae, we used glume, lemma, palea or lodicule length. For Cyperaceae, we used glume length. For dimorphic taxa (e.g. Rubiaceae, genus *Coprosma*) we characterized both male and female flowers.

To test whether the evolutionary trajectory of island flowers mirrors the patterns of their putatively correlated traits, we measured leaf area and seed dry mass for all species. These two traits were selected because, among the studied traits of island plants, they are the ones most clearly under direct selective pressures (Kavanagh and Burns, 2014; Biddick et al., 2019; Whittaker et al., 2023; Ciarle et al., 2024). Leaf shape was approximated as an ellipse so that leaf area was calculated from leaf length and width using the equation:

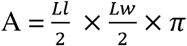

where A is the estimated leaf area, Ll is leaf length and Lw is leaf width. Seed mass was estimated from seed length using the equation:

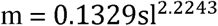

where m is the estimated seed mass, sl is the seed length and the two numeric values are constants. The equation was parametrized based on a linear model with both variables log-transformed, a sample size of 695 species with directly measured seed mass (74% of which are native species) across 297 genera and 100 families, and a fit of R squared = 0.865 (Sarah Richardson, personal communication, May 20, 2020).

All traits were extracted from the Floras of New Zealand and Australia series (Allan, 1961; Moore and Edgar, 1970; Webb et al., 1988; Edgar et al., 2010) and additional online sources (New Zealand Plant Conservation Network, 2023; Atlas of Living Australia, 2023; and New South Wales Flora Online, 2023). We searched the primary literature when data were missing. When online data was unavailable, we manually measured floral traits from herbaria in the Botany collection of Te Papa Tongarewa Museum of New Zealand. For each species trait, 5 to 10 specimens with multiple, well-preserved flowers were sampled. Three to five measurements were then taken for each specimen and averaged. When possible, to take into account temporal and spatial variability, specimens of different ages and from different regions of New Zealand were selected for each species.

### Statistical analyses

In this study we used two types of trait data: herbaria measurements and data retrieved from other sources. To test for significant differences between data types, we randomly selected 30 species for which data was available online and measured them in the herbarium collection. These data were then compared using a Wilcoxon signed-rank test.

For all analyses, we used the same trait for all species within a family (e.g. capitulum diameter for all Asteraceae). When a trait value was missing in a species, we used another trait in its place. To make sure this did not alter the results significantly, we tested whether the replacing trait correlated with the original one for all species of that family by using a Pearson’s correlation test (Berry and Feldman, 1985). A cut-off value of 0.7 was chosen, and only traits above the cut-off value were considered as possible replacements (Table S4).

To test whether floral display (i.e. flower size ⋅ number of flowers per inflorescence) of island species increased/decreased in size, we computed a floral display log ratio (fdLR) for each island/mainland pair of animal-pollinated plants as log(mean island floral display/mean mainland floral display). We then used a one-sided t-test to test whether fdLR was significantly above or below zero.

To test whether flower size of both animal- and wind-pollinated species followed the island rule, we computed a flower size log ratio (fsLR) as log(mean island flower size/mean mainland flower size). We then conducted two major axis regressions using fsLR as the dependent variable and log(mean mainland flower size) as the independent variable. Slopes significantly lower than zero would be evidence for the island rule. Slopes indistinguishable from zero would denote morphological isometry (Lomolino et al., 2013). Intercepts above or below zero would be evidence for gigantism or dwarfism respectively (Burns 2022). To avoid having the independent variable also as part of the dependent variable, the analysis was repeated without the fsLR ratio, with log(mean island flower size) as the dependent variable and log(mean mainland flower size) as the independent variable. Slopes significantly lower than one would be evidence for the island rule, while slopes indistinguishable from one would denote morphological isometry.

To test whether the evolutionary trajectories of flower size mirror the patterns of leaf area and seed mass, we computed a leaf area log ratio (laLR) as log(mean island leaf area/mean mainland leaf area), and a seed mas log ratio (smLR) as log(mean island seed mass/mean mainland seed mass). We then calculated Pearson’s correlations among fsLR, laLR and smLR for animal- and wind-pollinated species.

Mixed effect models were used to control for potential confounding factors. Because flower size may differ between plants with different breeding strategies, the breeding system was included as a fixed factor with two categories: monomorphic and dimorphic. Because flowers from different mainlands may respond differently to insular systems, mainland source was included as a fixed factor with two categories: Australia and New Zealand. Because we expect fully differentiated species to exhibit more pronounced insular trends compared to partially differentiated species (i.e. subspecies), taxonomic differentiation was included as a fixed effect with two categories: partially differentiated and fully differentiated. To control for phylogenetic conservatism, taxonomic families were included as a random effect. To avoid singularity, only taxonomic families with three or more island/mainland pairs were included. To control for environmental and ecological differences among islands, islands were included as a random effect. Since flower size may differ between different flower morphologies of animal-pollinated species, flower type was included as a fixed effect with three categories: actinomorphic flowers with fused petals; actinomorphic flowers with free petals, and zygomorphic flowers.

All analyses were repeated including only the island/mainland pairs identified through phylogenetic evidence to account for morphological uncertainty. All analyses were conducted in the R environment version 4.3.1 (R Core Team, 2022). Reduced major axis regressions were conducted using the R package *lmodel2* (Legendre and Oksanen, 2018). Mixed effect models were conducted using the R packages *nlme* (Pinhero et al., 2017) and *lme4* (Bates et al., 2009). When used as the independent variable, mainland flower size was log-transformed to avoid heteroscedasticity. *ggplot2* was used for data visualization (Wickham et al., 2016).

## Results

We identified 155 endemic island species. From these, we could compile island/mainland pairs for 145 species belonging to 43 families and representing 129 phylogenetically independent colonization events. We determined 85 island/mainland pairs based on phylogenetic evidence, comprising all representatives of a genus on the mainland. The remaining 44 island/mainland pairs were determined based on morphological evidence. We identified 90 animal-pollinated pairs and 39 wind-pollinated pairs. Floral data for 102 pairs was gathered from online sources, while data for the remaining 27 pairs was manually measured from herbaria specimens (Supplementary 1). Results of the Wilcoxon signed-rank test indicated no significant differences between herbarium measurements and data obtained from other sources (V = 177.5, P = 0.39). For 5 families, we retrieved data for multiple floral traits. Pearson’s correlation revealed all within-family traits to strongly correlate (Pearson’s correlation coefficient > 0.7, Supplementary 3, Table S1).

Floral display in animal-pollinated species exhibited no trend toward gigantism or dwarfism (n = 86, LR mean = 0.03, t = 0.89, P = 0.37) (Fig 2a). Similarly, floral display in wind-pollinated species exhibited no trend toward gigantism or dwarfism (n = 30, LR mean = 0.01, t = 0.13, P = 0.89) (Fig 2b). Flower number alone exhibited no trend as well (Supplementary 3, Table S2).

**Figure 2.**
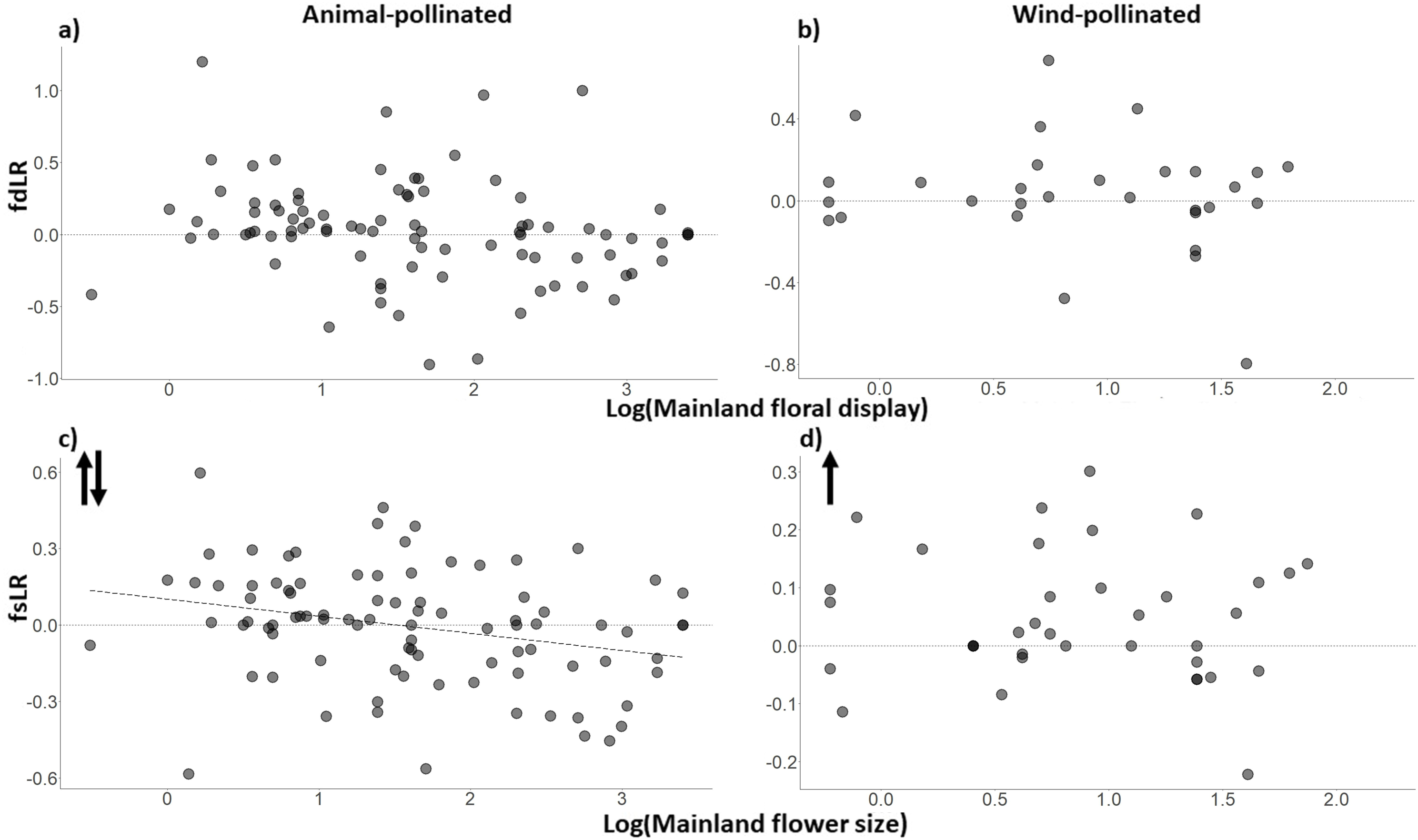
Floral display log ratio (fdLR) and flower size log ratio (fsLR) of animal- and wind-pollinated species (y-axis) varying as a function of log mainland values (x-axis) for a) animal-pollinated floral display, b) wind-pollinated floral display c) animal-pollinated flower size and d) wind-pollinated flower size. Each point denotes an island/mainland pairing. Floral display follows no recognisable pattern for species of either pollination mode. For animal-pollinated species, the flower size slope is significantly smaller than 0, giving support for the island rule. For wind-pollinated species, the flower size slope does not vary significantly from 0, but most points are above the horizontal line, indicating a trend toward gigantism. Opposing arrows denote the island rule. Single, upward arrows denote gigantism.

Flowers of animal-pollinated plants did not get larger (n = 87, LR mean = -0.01, t = -0.23, P = 0.81) but followed the island rule, with a slope lower than zero (s = -0.15, 95% confidence interval = -0.27, -0.04; P < 0.01) (Fig 2c). Instead, flowers of wind-pollinated species evolved toward gigantism (n = 36, LR mean = 0.05, t = 2.50, P = 0.01) and did not follow the island rule (s = -0.09, 95% confidence interval = -0.23, 0.06; P = 0.81) (Fig 2d).

These patterns did not mirror those of leaf area and seed mass. fsLR and laLR did not correlate for animal-(r = 0.21, P = 0.26) and wind-pollinated species (r = 0.17, P = 0.31). Similarly, fsLR and smLR did not correlate for animal-(r = 0.26, P = 0.12) and wind-pollinated species (r = 0.34, P = 0.07).

Results remained consistent when only the 85 island/mainland pairs identified through phylogenetic evidence were included in the analyses (Supplementary 3). Results did not vary when mean island flower size (log-transformed) was used as the dependent variable instead of fsLR. Results remained robust after controlling for breeding system, mainland source, degree of taxonomic differentiation, phylogenetic conservatism (i.e. taxonomic families), and islands (Supplementary 3, Table S3). In animal-pollinated species, fsLR did not vary significantly among mainland sources, taxonomic families, and degrees of taxonomic differentiation, but did so among breeding systems and islands (Supplementary 3, Table S4). In wind-pollinated species, fsLR did not vary significantly between breeding system, mainland source, degree of taxonomic differentiation, taxonomic families, and islands (Supplementary 3, Table S4).

## Discussion

We provided the first evidence for generalized *in-situ* evolution of flower size on islands and the first test for the island rule in flower size. Different pollination modes followed different evolutionary trajectories. We found no evidence for increased/decreased floral display in animal-pollinated species. Instead, flower size of animal-pollinated species followed the island rule. Conversely, flower size of wind-pollinated species became consistently larger on islands. However, neither of these patterns mirrored those of leaf area and seed mass. While the scale of our analysis did not allow us to determine which mechanisms produced the observed patterns, it is possible to suggest some mechanistic hypothesis based on our results and previous studies.

The floral display of animal-pollinated species exhibited no recognizable pattern, becoming neither larger nor smaller. To be advantageous, resources allocated to flowers have to increase reproductive success, which is pollinator-driven. Floral displays are also subjected to non-pollinator selective pressures such as herbivores and environmental stresses, which determine flower maintenance costs. If pollinators decline, but reproductive success still increases at a higher rate than maintenance costs, selection will favour increased pollinator attraction. Only when pollinators become too scarce and/or mating partners are limited will selection favour individuals that rely on abiotic vectors/self-fertilization (Maciel et al., 2020; Roddy et al., 2021). Consequently, depending on the biotic and abiotic factors of islands, pollinator paucity could result in showy flowers, inconspicuous flowers, or neither. Our results indicate that, if pollinators do decline on the islands of the Southwest Pacific, differences in pollinator pressures between islands and the mainland might be negligible and/or countered by differences in flower maintenance costs.

Flower size of animal-pollinated flowers followed the island rule. Flower size is known to correlate with other plant traits, including leaf area (Niklas, 1994; Burns et al., 2012; Bawa et al., 2019; Roddy et al., 2021; ELVojtkó et al., 2022). However, results indicate that flower size of island species evolved independently of leaf area and seed mass. Thus, it is unlikely that selection acting on these traits caused correlated evolution in flower size. Flower size may also follow the island rule because of the relationship between the size of flowers and their respective pollinators (Burns, 2019). Size-coupling is a key aspect of plant-pollinator mutualism in specialized flowers (Dalsgaard et al., 2009; Maglianesi et al., 2014; Biddick & Burn, 2018). Consequently, if the smallest and largest pollinator species were excluded from island colonization, this could drive the smallest and largest flower species to shift toward intermediate sizes, conforming to the island rule (Hiraiwa and Ushimaru, 2017; Burns, 2019). However, this is unlikely to apply to our system as most of our island endemics possessed open, radial, generalist flowers that would likely not comply with size changes in pollinators (Faure et al., 2022). It is also relevant to note that pollinator specialists are rare on most islands (Olesen et al., 2002; Traveset et al., 2016; Wang et al., 2020; Whittaker et al., 2023). Biddick and Burns (2021) showed how a null model only accounting for evolutionary drift can provide a parsimonious explanation for the island rule. As a mechanistic explanation for why some plant traits obey the island rule and others do not is still lacking (Biddick et al., 2019; Whittaker et al., 2023), it could be beneficial to look outside mechanisms of selection to explain why flower size of animal-pollinated species follows the island rule.

Results for animal-pollinated species varied significantly between breeding systems (Figure S1). Specifically, flowers of dimorphic animal-pollinated species became larger, while flowers of monomorphic animal-pollinated species did not (Supplementary 3, Fig S1a). Globally, pollinator selection acts more strongly on dimorphic species, which usually possess small flowers (Geber et al., 2012). Evolution toward larger flowers in males of dimorphic animal-pollinated island plants may therefore reflect evolution in a depauperate environment where every possibility of attracting pollinators needs to be exploited (Godley, 1979). Additionally, differences in breeding systems between animal-and wind-pollinated species might be explained by considering that, while females allocate more to reproduction in dimorphic animal-pollinated plants, in wind-pollinated species the reproductive costs of males may exceed or match those of females, potentially causing plants with different pollination modes to follow different evolutionary trajectories (Harris and Pannell, 2008; Liu et al., 2021; Roddy et al., 2021; but see Midgley, 2022).

Flower size of wind-pollinated species did not follow the island rule but evolved to be consistently larger on islands. As with animal-pollinated species, results indicate that flower size evolved independently of leaf area and seed mass. Gigantism in wind-pollinated flowers could be explained as a by-product of reduced dispersibility. In wind-dispersed species, smaller pollen usually has a higher dispersal ability, which on islands increases the chances of being blown out at sea (Burns, 2019). Island conditions may therefore select for larger pollen, which is a trait that can correlate with larger flowers (Torres, 2000; Anguilar et al., 2002). Flower gigantism could also result from loss of defence. The loss of defence mechanisms is a hallmark of oceanic island plants (Ciarle et al., 2024), and a decrease in resources allocated to defence could allow an increase in resources allocated to reproduction. Finally, larger flowers could correlate with larger stigmas capable of more effective pollen-capture, thus leading to increased pollination success (but see Friedman and Barrett, 2009; Cresswell et al., 2010).

However, why wind-pollinated flowers would evolve toward gigantism on islands is also puzzling. First, wind-pollinated plants typically benefit from producing many small, uniovulate, inexpensive flowers. This is because numerous flowers can sample more of the airstream and maximize pollen capture and seed production (Sakai and Sakai, 1995; Friedman and Barrett, 2011). According to this model, producing larger, more expensive flowers would reduce maternal fitness, and possessing larger flowers could be beneficial only if island plants possessed more resources available for flower and seed production than their mainland relatives.

Second, seed gigantism is a hallmark of New Zealand’s outlying islands (Kavanagh and Burns, 2014; Biddick et al., 2019) and previous studies on continents indicate that larger seeds often correlated with smaller flowers (Bawa et al., 2019). Larger seeds can lead to increased within-fruit sibling competition. One of the strategies adopted by plant species to counter sibling competition is to reduce the number of seeds per fruit (Bawa, 2016). While this potentially reduces fitness, it also reduces flower and pollinator investment, allowing plants to produce small flowers that are wind-pollinated or that rely on generalist pollinators (Bawa et al., 2019). In other words, as seed size increases, flower size of wind-pollinated plants should decline, which is not what we observe. Again, perhaps insular wind-pollinated plants allocate resources differently than their mainland counterparts, promoting consistent positive flower/seed correlations. The reason behind it though remains unclear.

The size of both animal- and wind-pollinated flowers showed predictable patterns while floral display did not. Floral display (i.e. flower number ⋅ flower size) quantifies the overall amount of resources allocated to flower production, while flower size can inform how these resources are allocated within individual reproductive structures. Observed differences may indicate that island and mainland species allocate the same amount of resources to flower production, but they do so differently. Specifically, wind-pollinated species on islands seem to invest into fewer and larger flowers, while animal-pollinated species appear to invest into fewer and larger or more and smaller flowers depending on their flower size at the time of island colonization.

Differences in pollination agents may cause wind- and animal-pollinated species to follow different evolutionary trajectories. For example, pollinator size-coupling could produce an island-rule-like pattern only in animal-pollinated species, while possessing larger stigmas would only be advantageous in wind-pollinated flowers. However, beyond pollination mode, the two groups seem rather heterogeneous. For example, they both contain animal- and wind-dispersed species spanning multiple plant families, with several families featuring in both groups. This difference may just be idiosyncratic to our system. We encourage future studies to conduct similar analysis on other island systems, and to design small case studies capable of discerning different possible mechanisms.

Our analysis encompasses only a portion of the Southwest Pacific. First, the study does not include any tropical islands. Ecologically, tropical systems might operate differently than non-tropical ones. For instance, while pollination networks seem to be structurally similar in tropical and non-tropical areas (Vizentin-Bugoni et al., 2018), many plant traits follow a latitudinal gradient, including flower morphological diversity (Currie and Paquin, 1987; Chen et al., 2017; Chartier et al., 2021). Additionally, tropical systems are underrepresented in the literature (Culumber et al., 2019). Secondly, our study focuses on small, isolated archipelagos with relatively small species pools. As the ecological and evolutionary dynamics of islands vary markedly with area and isolation (MacArthur and Wilson, 1967; Whittaker et al., 2023), these results may not apply to larger and less isolated archipelagos.

In this study, we accounted for phylogenetic relatedness through paired taxonomic comparisons and by adding taxonomic family as a random effect in our mixed models. To make sure that flower size evolution was widespread among flowering plants we included multiple families and two mainland source pools. However, accounting for relatedness using a systematic approach, such as a phylogenetic generalized least square model, would have likely yielded more accurate results. This was not possible as a well-resolved phylogeny for the investigated species is not currently available.

Our focus was on many island/mainland pairs of species, spanning several families. However, many families in the analysis were only represented by one or a few species, making it difficult to evaluate within family trends. Moreover, focusing on island/mainland pairs implies that we did not account for intraspecific variance. It is possible that accounting for such variance would alter the observed patterns. Finally, while our study focused on flower size, future research should aim for a multi-trait approach capable of encompassing the plant island syndrome in its entirety.

In conclusion, we provided the first test for the island rule in flowers, and the first evidence for generalized *in-situ* evolution of flower size on islands. Results indicate that, while *in situ* evolution of flower size is widespread on islands in the Southwest Pacific, species belonging to different pollination modes exhibit markedly different evolutionary trajectories. These results combine with a growing body of literature investigating the island syndrome in plants (Burns, 2019; Schrader et al., 2021; Delavaux et al., 2021; Meudt et al., 2021; Zizka et al., 2022; Whittaker et al., 2023; Ciarle, 2024), and provide a much-needed insight into the evolution of plants on islands.

## Acknowledgments

### Acknowledgements

We thank Bridget Hatton for helping us navigate the Kaitiaki Taonga Herbarium Collection at Te Papa Tongarewa Museum of New Zealand. We thank Alessandra Franco for drawing Figure 1. We also thank Sarah Richardson for sharing her data on seed mass.

## Ethics

This work did not require ethical approval from a human subject or animal welfare committee.

## Data availability Statement

Plant species lists, flower traits measured, and models used for the analysis are available in the supplementary materials of this article.

## Supplementary data

Supplementary 1: List of Island-Mainland pairs and sources used as references.

Supplementary 2: List of floral traits and data used for the analysis.

Supplementary 3: Supplementary tables and figures.

