## Supplementary 3 for "Flower size evolution in the Southwest Pacific"

**Reduced major axis regression (RMA)**

Using LR in the regression models implies that mainland flower size is included as both dependent and independent variable. To avoid this, we repeated the RMA analysis using island flower size as the dependent variables and mainland flower size as the independent variables. Results remained consistent. Animal-pollinated flowers followed the island rule, with a slope smaller than 1 (s = 0.84, 95% confidence interval = 0.73, 0.96; P < 0.001). Wind-pollinated flowers did not follow the island rule (s = 1.01, 95% confidence interval = 0.86, 1.19; P < 0.001).

**Analyses with only phylogenetically-identified pairs**

To remove morphological uncertainties from the results, analyses were repeated using only the 85 island/mainland pairs identified through phylogenetic evidence. Results remained consistent. Floral display in animal-pollinated species exhibited no trend toward gigantism or dwarfism (n = 59, LR mean = 0.56, t = 1.34, P = 0.19). Similarly, floral display in wind-pollinated species exhibited no trend toward gigantism or dwarfism (n = 15, LR mean = 0.19, t = 0.21, P = 0.84). Flowers of animal-pollinated plants did not get larger (n = 59, LR mean = 0.06, t = 1.34, P = 0.19) but followed the island rule, with a slope lower than zero (s = -0.31, 95% confidence interval = -057, -0.10; P < 0.01). Instead, flowers of wind-pollinated species evolved toward gigantism (n = 20, LR mean = 0.06, t = 2.27, P = 0.03) and did not follow the island rule (s = -0.15, 95% confidence interval = -082, 0.33; P = 0.81).

**Model selection**

The RMA used to test the island rule in animal-pollinated and wind-pollinated species was: LR ~ intercept + LM + error. When building the mixed models, because different islands had different subsets of taxonomic families, families and islands could not be included in the same model. We started by fitting four separate models, two for animal-pollinated flowers and two for wind-pollinated flowers.

- For animal-pollinated flowers:
  I. LR ~ intercept + LM * Flower morphology * Breeding System * Mainland source + (island) + error
  II. LR ~ intercept + LM * Flower morphology * Breeding System * Mainland source + (Family) + error
- For wind-pollinated flowers:
  I. LR ~ intercept + LM * Breeding System * Mainland source + (island) + error
  II. LR ~ intercept + LM * Breeding System * Mainland source + (Family) + error.

In all models, the random effect is shown in brackets. LM indicates log(Mainland flower size). LR is log(Island flower size/Mainland flower size). We then proceeded with model selection, first by finding the optimal structure of the random component using the AICc criterion, and then by finding the optimal fixed structure by looking at the significance of each fixed component. Finally, we validated the models by checking for normality of the residuals and violation of homogeneity. Best models are reported in Table S3.

**Differences between breeding systems**

Mixed model results showed a significant difference between monomorphic and dimorphic species of animal-pollinated flowers (Fig S1a). This was confirmed by further analysis indicating that flowers of dimorphic animal-pollinated species get larger (n = 25, LR mean = 0.87, t = 2.81, P = 0.001), while flowers of monomorphic animal-pollinated species do not (n = 56, LR mean = -0.04, t = -1.53, P = 0.13).

**Table S1** Pearson’s correlation coefficients of flower traits across families for which multiple traits were measured. Columns indicate the family, the two traits correlated in each instance and the Pearson’s correlation coefficient. Cut-off value was set at 0.7 (Berry & Feldman, 1985).

| Family | Traits | Coefficient |
| --- | --- | --- |
| Boraginaceae | Lobe length~Flower diameter | 0.92 |
| Asteraceae | Disk diameter~Total diameter | 0.89 |
| Asteraceae | Ray floret length~Total diameter | 0.84 |
| Asteraceae | Disk diameter~Ray floret length | 0.75 |
| Gentianaceae | Tube width~Lobe length | 0.98 |
| Poaceae | Lodicules length~Lemma length | 0.94 |
| Poaceae | Lodicules length~Palea length | 0.93 |
| Poaceae | Lodicules length~Glumes length | 0.80 |
| Poaceae | Lemma length~Glumes length | 0.82 |
| Poaceae | Lemma length~Palea length | 0.91 |
| Poaceae | Palea length~Glumes length | 0.66 |
| Rubiaceae | Male corolla length~Female corolla length | 0.65 |

**Table S2** One-sided t-tests for flower numbers. No pattern toward gigantism or dwarfism can be detected for animal- and wind-pollinated flowers.

| Group | n | LR mean | t | P |
| --- | --- | --- | --- | --- |
| Animal-pollinated | 80 | 0.09 | 1.43 | 0.16 |
| Wind-pollinated | 31 | -0.04 | -0.52 | 0.61 |

**Table S3** Models selected for animal-pollinated and wind-pollinated flower size. The first model is the one initially used to test the island rule. The additional models are the best models selected after all confounding factors were added as fixed or random effects. Only one model is presented for wind-pollinated flowers as the initial RMA model was also selected as the best model. The estimated slope is presented along with the 95% confidence intervals (in squared brackets). Significant p-values (< 0.05) indicate negative slopes and are evidence for the island rule. LR = log(Island flower size/Mainland flower size). LM = log(Mainland flower size). Significant p-values are highlighted with ** (P < 0.05).

|  | Model | Slope | P |
| --- | --- | --- | --- |
|  | LR ~ intercept + LM + error | -0.15 [-0.45, -0.09] | 0.01** |
| Animal-pollinated flowers | LR ~ intercept + LM + (island) + error | -0.12 [-0.19, -0.01] | 0.02** |
|  | LR ~ intercept + LM + Breeding system + (Family) + error | -0.08 [-0.13, -0.01] | 0.04** |
| Wind-pollinated flowers | LR ~ intercept + LM + error | -0.01 [-0.13, 0.10] | 0.81 |

**Table S4** Results of Kruskal-Wallis analysis of variance, used to assess significant differences in fsLR means among breeding systems, mainland sources, degree of taxonomic differentiation, taxonomic families and islands. Significant differences are highlighted with ** (P < 0.05).

|  | Animal-pollinated flowers | Wind-pollinated flowers |
| --- | --- | --- |
| Breeding system | Chi-squared = 6.12, df = 1, P = 0.04** | Chi-squared = 0.43, df = 1, P = 0.14 |
| Mainland source | Chi-squared = 2.57, df = 1, P = 0.23 | Chi-squared = 2.25, df = 1, P = 0.14 |
| Degree of taxonomic differentiation | Chi-squared = 2.04, df = 1, P = 0.34 | Chi-squared = 0.39, df = 1, P = 0.22 |
| Phylogenetic conservatism (Taxonomic families) | Chi-squared = 9.10, df = 8, P = 0.18 | Chi-squared = 0.74, df = 3, P = 0.14 |
| Islands | Chi-squared = 18.98, df = 10, P = 0.04** | Chi-squared = 6.37, df = 6, P = 0.39 |

**Figure S1** Differences in fsLR means among a) breeding system, b) mainland, c) degree of taxonomic differentiation, d) taxonomic families and e) islands. Families with less than four island/mainland pairs are not shown. Snares and Macquarie islands are excluded as each island only housed a single endemic species.

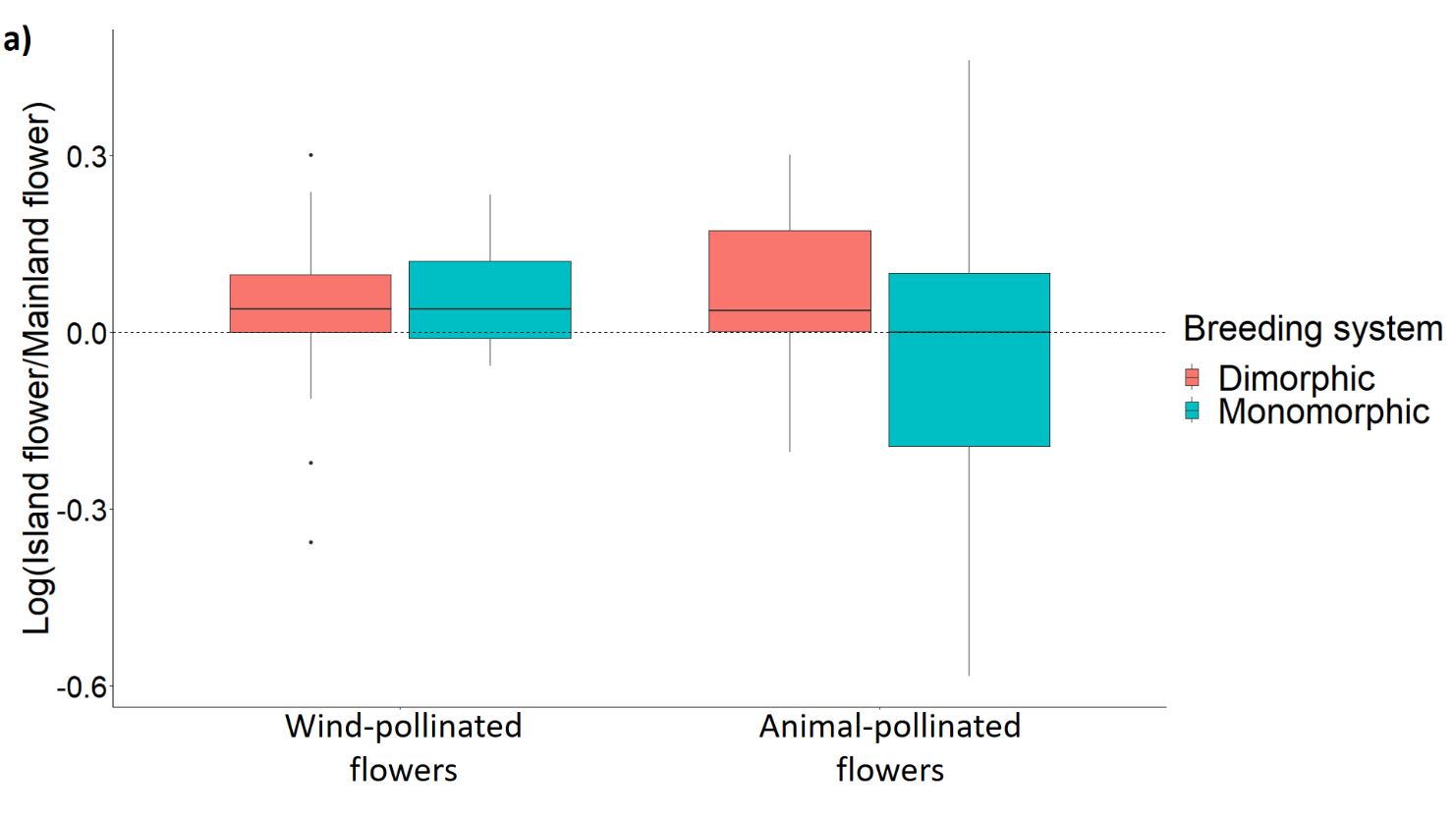

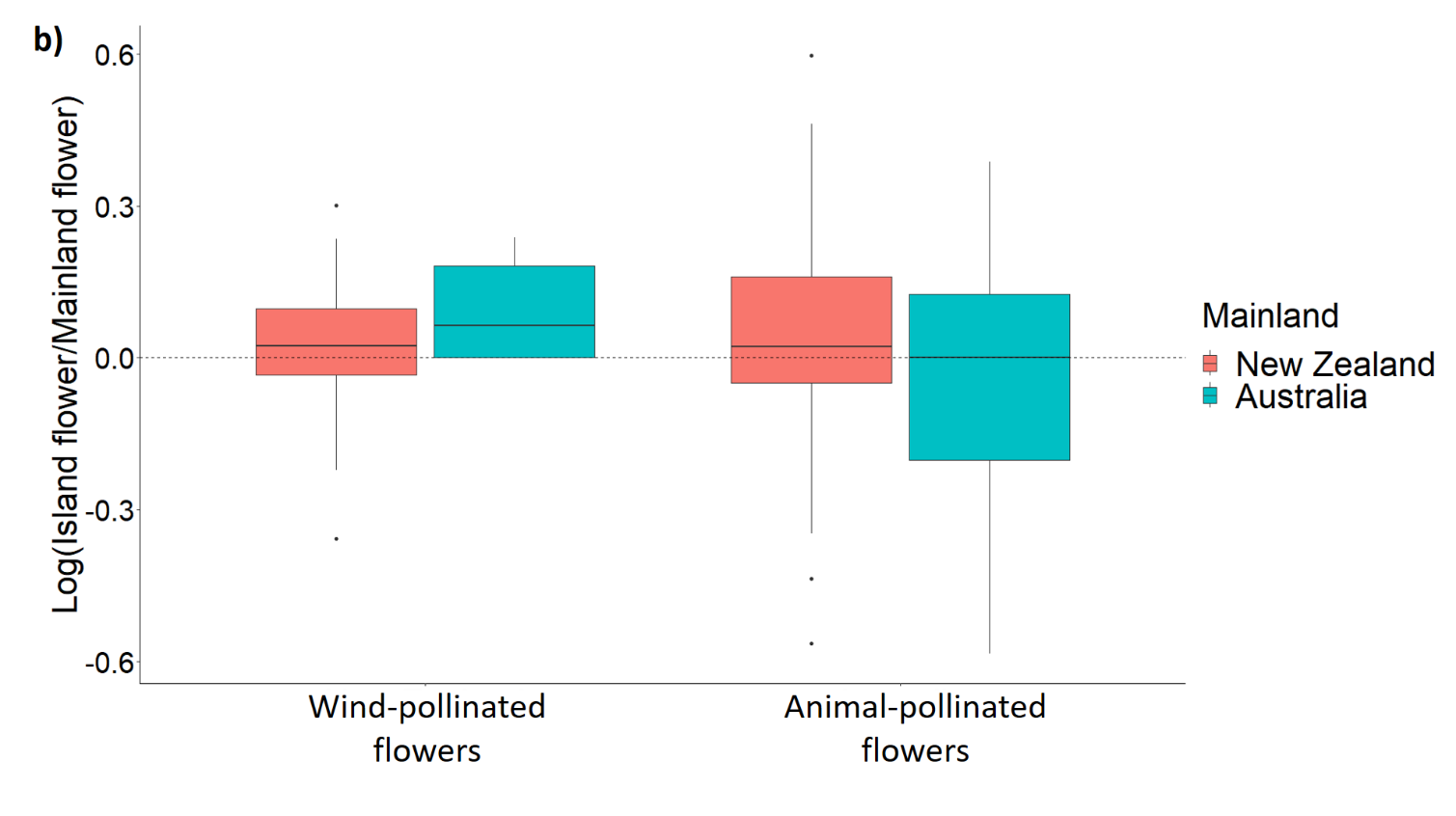

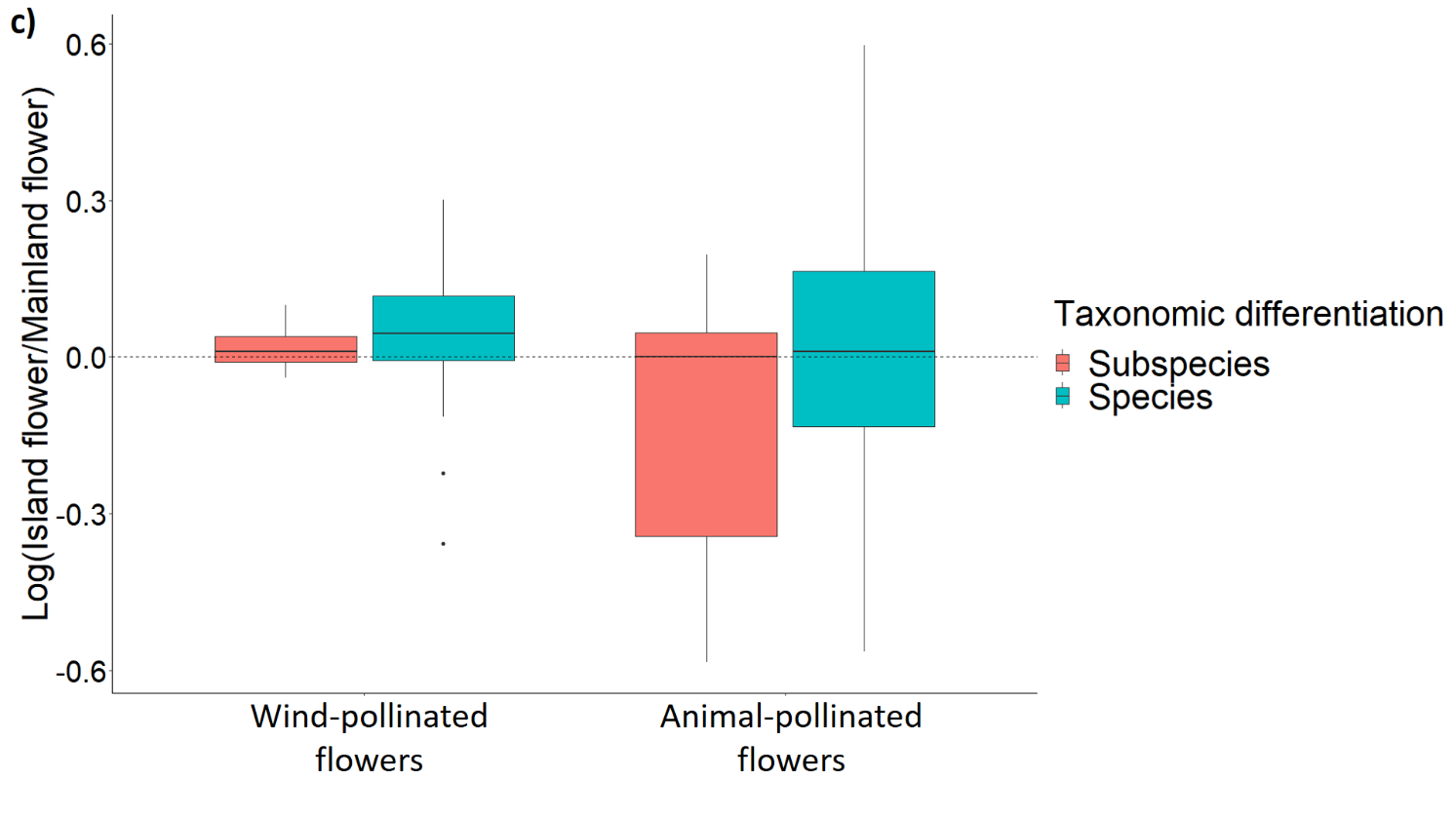

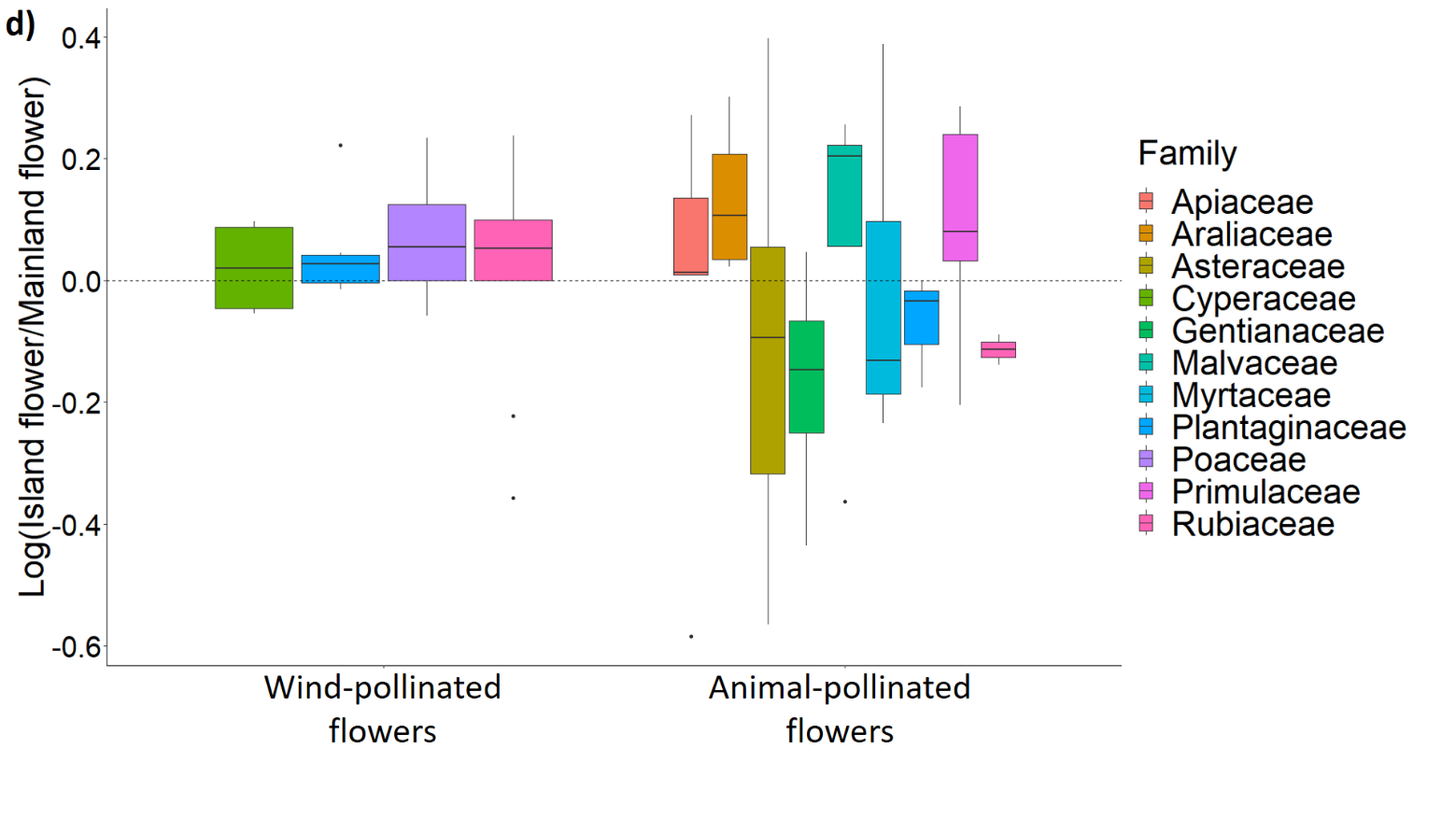

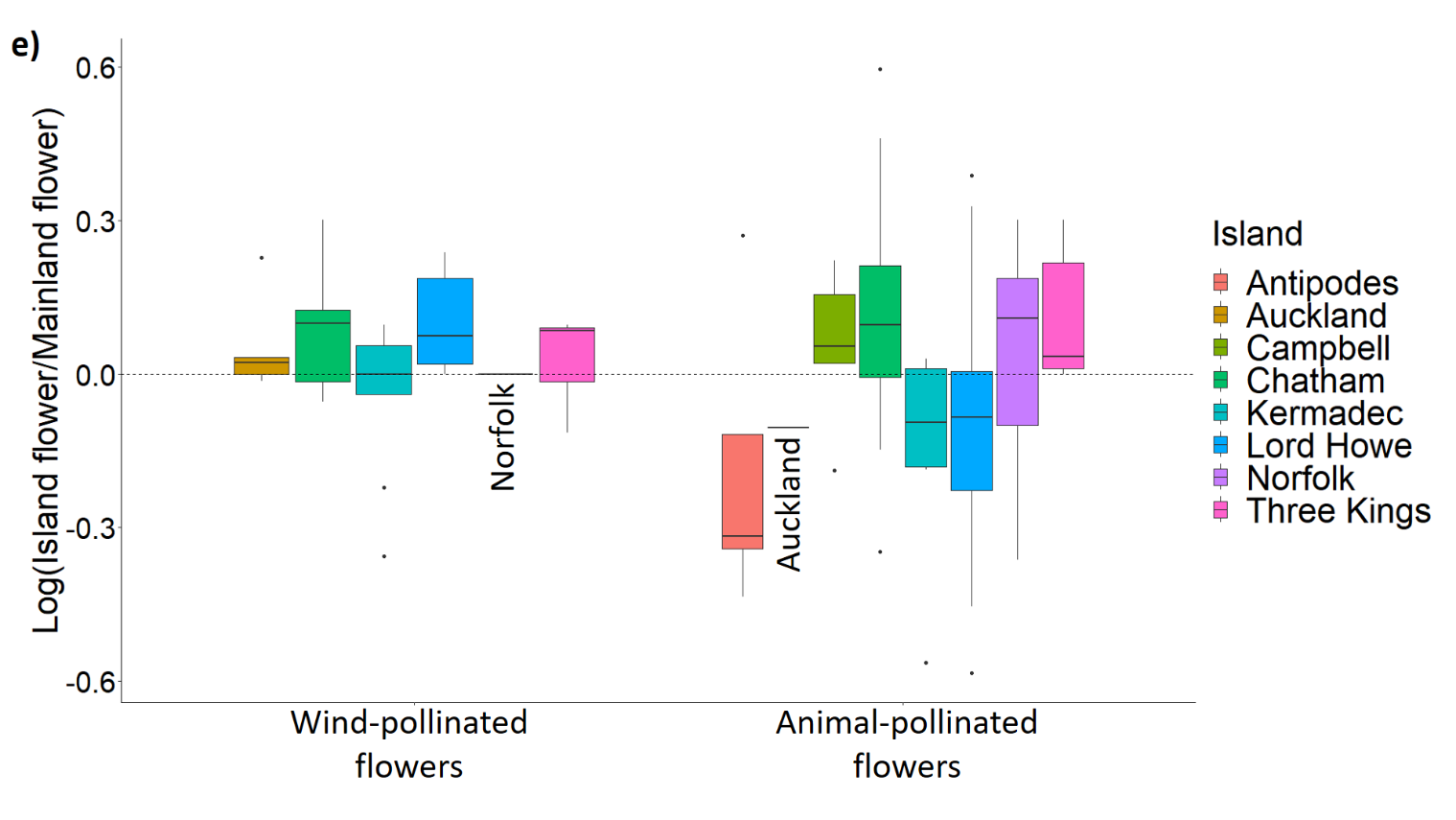
